# Effect of vitamin D supplementation on biomarkers of inflammation and immune function: functional genomics analysis of the BEST-D trial

**DOI:** 10.1101/217612

**Authors:** Antonio J. Berlanga-Taylor, Katherine Plant, Andrew Dahl, Evelyn Lau, Michael Hill, David Sims, Andreas Heger, Jonathan Emberson, Jane Armitage, Robert Clarke, Julian C. Knight

## Abstract

Vitamin D deficiency has been associated with multiple diseases, but the causal relevance and underlying processes are not fully understood. Elucidating the mechanisms of action of drug treatments in humans is challenging, but application of functional genomic approaches in randomised trials may afford an opportunity to systematically assess molecular responses to treatments. In the Biochemical Efficacy and Safety Trial of Vitamin D (BEST-D), 305 community-dwelling individuals aged over 65 years were randomly allocated to treatment with vitamin D_3_4000 IU, 2000 IU or placebo daily for 12 months. Genome-wide genotypes at baseline, and transcriptome and plasma levels of cytokines (IFN-γ, IL-10, IL-8, IL-6 and TNF-α) at baseline and after 12 months, were measured. The trial had >90% power to detect a 2-fold change in gene expression. Allocation to vitamin D for 12-months was associated with 2-fold higher plasma levels of 25-hydroxy-vitamin D (25[OH]D), but had no significant effect on whole-blood gene expression (FDR <5%) or on plasma levels of cytokines compared with placebo. In pre-specified analysis, rs7041 (intron variant, *GC*) had a significant effect on circulating levels of 25(OH)D in the low dose but not on the placebo or high dose vitamin D regimen. A gene expression quantitative trait locus analysis (eQTL) demonstrated evidence of 31,568 *cis*-eQTLs (unique SNP-probe pairs) among individuals at baseline and 34,254 after supplementation for 12 months (any dose), but had no significant effect on cis-eQTLs specific to vitamin D supplementation. The trial demonstrates the feasibility of application of functional genomics approaches in randomised trials to assess the effects of vitamin D on immune function.

**One sentence summary:** Supplementation with high-dose vitamin D in older people for 12 months in a randomised, placebo-controlled trial had no significant effect on gene expression or on plasma concentrations of cytokines.

**Trial registration:** SRCTN registry (Number 07034656) and the European Clinical Trials Database (EudraCT Number 2011-005763-24).

**Funding:** Medical Research Council, British Heart Foundation, Wellcome Trust, European Research Council and Clinical Trial Service Unit, Nuffield Department of Population Health, University of Oxford, Oxford, United Kingdom

**Copyright:** Open access article under the terms of CC BY.

## Introduction

Randomised controlled trials afford reliable approaches to understand the causal relevance of drug treatments and can also help our understanding of disease mechanisms by relating changes in biomarkers with incidence of disease or with surrogate markers of disease. Advances in molecular methods now permit use of high-throughput functional genomics strategies in clinical trials. Application of such approaches has been under-utilised to date, with previous studies focusing on comparisons of transcriptomes to understand mechanisms and identify novel biomarkers (1). Animal and experimental models of disease pathogenesis have limited ability for translation into humans, particularly in the context of complex diseases (2, 3), while incomplete knowledge of mechanisms contributes to inconclusive findings in randomised trials and the current high failure rate in late stage drug development. Hence, there is an urgent need to demonstrate the likely significant value in combining functional genomic approaches such as genome-wide genotyping and gene expression profiling together with biochemical and clinical markers in clinical trials to enhance our understanding of pathophysiological processes and mechanisms of action of novel drug treatments. Such approaches may also allow high-throughput assessment of cellular and molecular responses at group and individual levels that can also be integrated with effects on clinical outcomes data. Hence, integrated analysis may yield clinically relevant insights about treatment that could guide the design of large outcome trials.

This study applied functional genomics methods to investigate the molecular response to vitamin D supplementation. In addition to the established role of vitamin D in calcium metabolism and bone disease, accumulating evidence suggests a possible role of vitamin D in immune function and inflammatory diseases (4). Previous studies have investigated the associations of vitamin D with gene expression (5-7), but these have typically been cross-sectional, in experimental models, involved relatively small sample sizes or lacked placebo controls. Moreover, no previous studies have assessed the impact of genetic variation on responses to vitamin D supplementation.

The aim of the work described here was to examine the molecular responses to vitamin D supplementation in a randomised, placebo-controlled trial. To achieve this, we investigated changes in response to treatment after 12 months in whole blood transcriptomes and plasma levels of cytokines, in addition to genetic determinants of individual responses on circulating 25-hydroxy vitamin D (25[OH]D) and genome-wide gene expression, by comparing a total of 305 individuals allocated to daily treatment with vitamin D at either 4000 IU, 2000 IU or placebo in the BEST-D trial (8).

## Results

### Effects of vitamin D supplementation on gene expression

Following sample processing and quality control, genome-wide gene expression data on 16,760 probes (12,910 genes) were available for 298 of 305 participants who were randomised to the trial (560 samples, of which 262 had both baseline and 12 month samples) (Figure 1). The mean age of study participants was 72 years at randomisation, 51% were male and 12% reported prior use of vitamin D supplements (≤400 IU of vitamin D_3_ daily) (9). Compliance with instructions to take vitamin D supplements or placebo was high, with 90% (4000 IU), 92% (2000 IU) and 85% (placebo) reporting taking the capsules on all or most days at 12 months (8, 9). The overall mean plasma level of 25(OH)D was 50 nmol/L (standard error [SE] 1.04) at baseline and treatment was associated with mean plasma levels of 25(OH)D of 136 (3.94). 106 (2.55) and 50 (1.68) among those allocated to 4000 IU, 2000 IU, and placebo, after 12 months of treatment (unadjusted levels) (Table 1).

**Figure 1:**
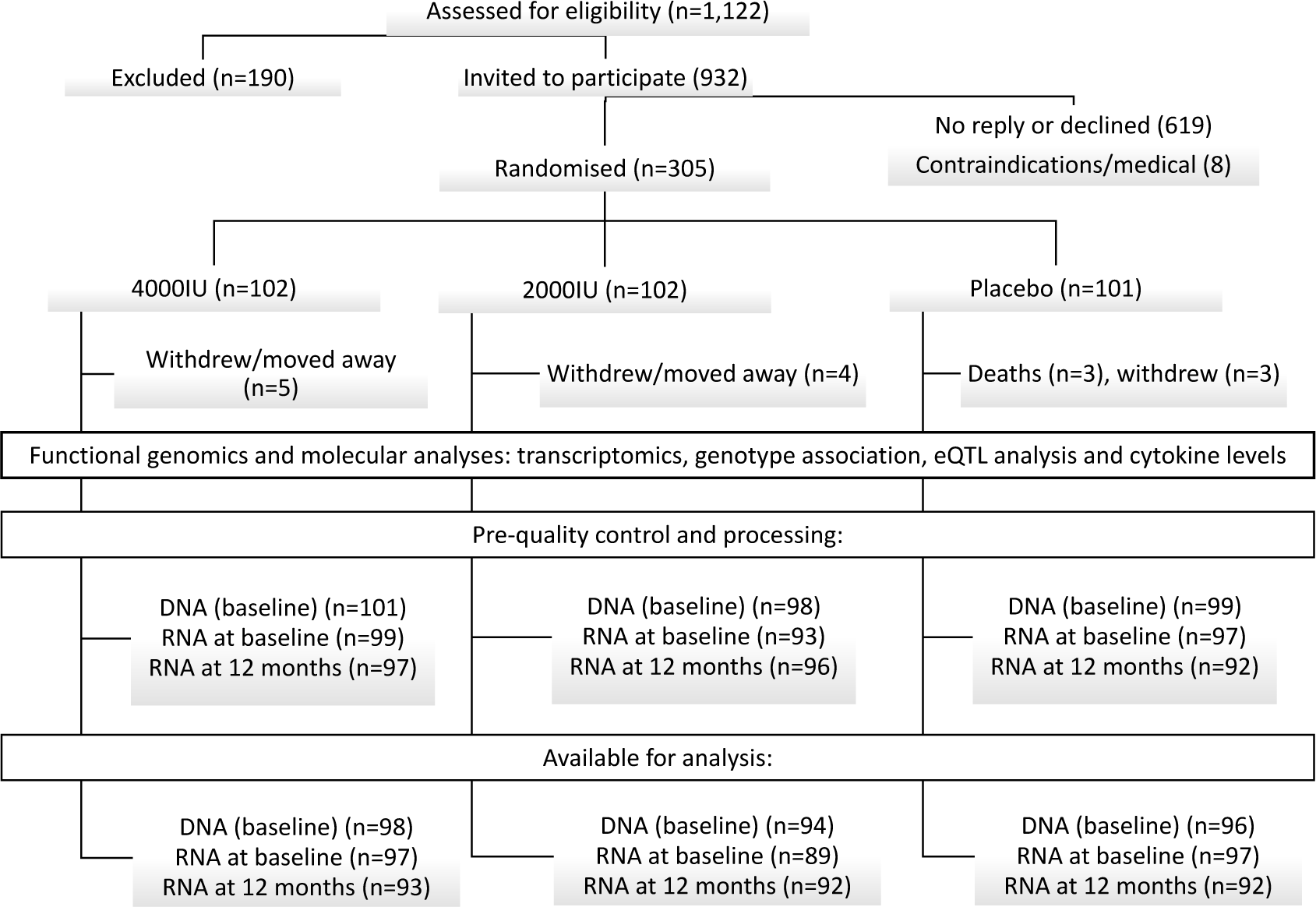
CONSORT diagram for the functional genomics analysis of participants in the BEST-D study. eQTL: expression quantitative trait loci.

We first performed principal components (PC) analysis to visualise the relationship between samples based on gene expression values. Considering all samples or paired samples within treatment or placebo groups, we found no visual evidence of clustering using up to the first 13 principal components (accounting for 41% of variance) (Figure 2, Supplementary Figure 2A and Supplementary Figures 3-6). The top 100 PCs accounted for 60% of the total variance (Supplementary Figure 3).

**Figure 2:**
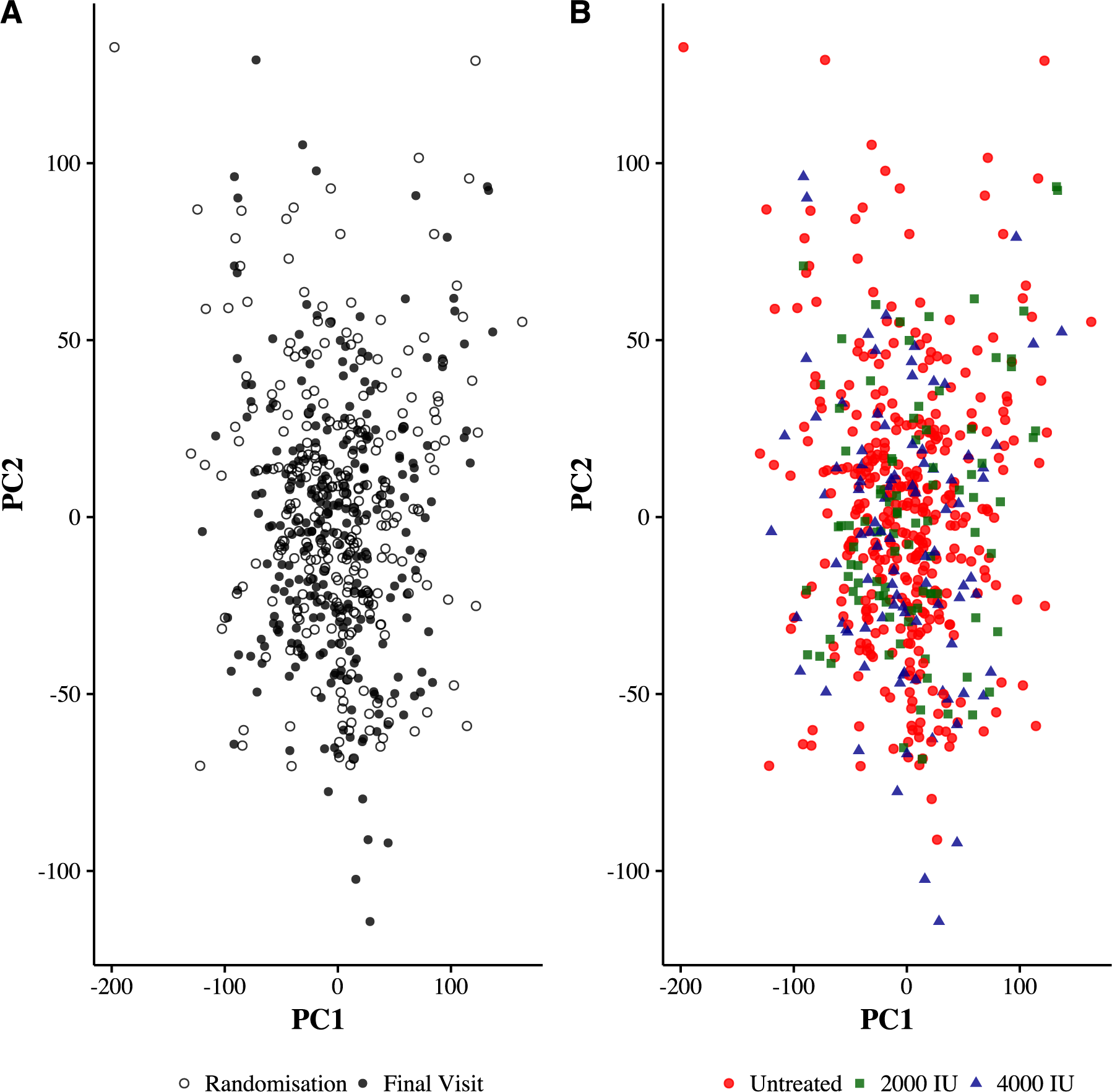
Global gene expression by allocated treatment with either vitamin D or placebo supplementation was assessed using principal components analysis of variance. Principal components (PC) were calculated based on post-QC and VSN normalised gene expression signals. PC1 and PC2 (representing 15% and 8.5% of variance, respectively) are shown and coloured according to time-point (final visit at 12 months or at randomisation [baseline], panel A) and separately by treatment group (placebo, 2000 IU or 4000 IU, panel B).

We next formally tested the effects of vitamin D supplementation on genome-wide gene expression considering significantly differentially expressed probes at an FDR <5% for any fold change in an unadjusted model. As expected after successful randomisation, we did not observe any significant differences in gene expression when comparing allocation groups at baseline (placebo baseline, 2000 IU baseline and 4000 IU baseline) (Supplementary Table 1).

The pre-defined primary outcome sought to determine differences in gene expression in response to any dose of vitamin D compared with placebo. We compared differences in transcriptome among participants allocated to vitamin D (4000 IU or 2000 IU) at 0 vs 12 months versus those allocated to placebo. Difference in difference analysis was estimated using (expression gene A following vitD_12months_ - expression gene A vitD_baseline_) - (expression gene A placebo_12months_ - expression gene A placebo_baseline_). We found that 375 probes (for 4000 IU minus placebo using the difference in difference estimate as above) and 329 probes (for 2000 IU minus placebo) were significantly differentially expressed following vitamin D supplementation (unadjusted p-value <0.05), but none remained significant after taking account of multiple comparisons (FDR <5%) (Supplementary Table 1). Use of quantile or variance stabilizing normalization (VSN) methods for data processing did not materially alter the results (Supplementary Figure 7, see Methods for details). The placebo group, with both baseline and 12 month sampling, is a stringent control that was used to account for the effect of time, placebo itself, technical artefacts or other sources of variation. After controlling for relevant confounders, there were no differences in gene expression between patients allocated vitamin D versus those allocated placebo. It is possible that reliable detection of gene expression in response to vitamin D supplementation may require a larger sample size (Supplementary Figure 7).

Subsequent analyses considered less conservative comparisons as pre-specified (see data analysis protocol, Supplementary Appendix). We conducted two group comparisons of each arm (4000 IU at 12 months vs 4000 IU at baseline; and separately, 2000 IU at 12 months vs 2000 IU at baseline), but did not find significant differences in gene expression (FDR <5%). We also compared 12-month samples (4000 IU at 12 months vs placebo at 12 months; 2000 IU at 12 months vs placebo at 12 months), but did not observe any significant differences (FDR <5%). Likewise, paired analysis for each of these comparisons did not yield significant differences (Supplementary Table 1).

To maximise power, we combined all vitamin-D allocated individuals before and after treatment (2000 IU plus 4000 IU vs their baseline samples, unpaired), but did not detect significant differences in expressed probes (FDR <5%). Paired sample comparisons (n = 186, joint 2000 IU and 4000 IU) for this grouping indicated some significant differences following vitamin D supplementation (143 probes at <5% FDR; 292 probes at FDR <10%, fold change range 0.83–1.12, Supplementary Table 1). However, neither of these analyses took account of the differences in the placebo group and, hence, were less robust than the analyses of differences and random-effects detailed above. These results would thus require validation in larger studies with similar design.

We hypothesised that differences in transcriptomes would be more evident when comparing individuals with low plasma levels of 25(OH)D. We compared those with pre-treatment plasma levels of 25(OH)D <50 nmol/L versus those with 25(OH)D >50 nmol/L, (124 vs 159, respectively, unadjusted model) regardless of the allocated treatment, and separately cases of more extreme change (<25 nmol/L vs >75 nmol/L 25(OH)D, 13 vs 30, respectively). Likewise, allocation to vitamin D had no significant differences in gene expression (FDR <5%, Supplementary Table 1). We repeated such comparisons using subsets with paired samples before and after treatment among individuals who at baseline were deficient (<50 nmol/L and separately for those with <25 nmol/L), but did not detect any significant differences in either subgroup (FDR <5%, Supplementary Table 1). Finally, although we did not pre-specify it, we selected individuals whose change (delta) in vitamin D levels was high (vitD_12months_ - vitD_baseline_), regardless of allocated treatment. Although this analysis is more likely to be confounded it provided more power as more individuals with greater differences in plasma vitamin D in response to supplementation were included. We chose the median (+44.79 nmol/L) as this yielded the highest change in the maximum number of individuals (n=145). As expected, none of the placebo group had a difference of this magnitude (median: +2.58 nmol/L). In paired analysis we found five genes significantly different at an FDR of <5% (Supplementary Table 1), but the effect sizes for these genes were small (fold change range: 0.81 – 1.12) and further work is needed to replicate such associations in other trial populations.

### Individual responsiveness to vitamin D supplementation: impact of genotype

This study lacked power to assess differences in the effects of treatment with vitamin D by differences in genome-wide genetic variation. We restricted our analysis to SNPs with prior evidence of association with plasma 25(OH) D levels from population GWAS (10) (rs12794714, rs2282679, rs7041 and rs7944926) for which we had genotyping data available. Previous studies in twins suggested that summer levels of vitamin D were not strongly influenced by genetic variation (11, 12). Our pre-specified analysis assessed the hypothesis that genotype may modulate the response to vitamin D supplementation. We analysed vitamin D levels following treatment with measurements at 6 and 12 months, using baseline vitamin D and other variables as covariates in a linear regression model (see Methods). We found that rs7041 (located on chromosome 4) was significantly associated with response to vitamin D treatment with low dose vitamin D (2000 IU) at 6 and 12 months (permuted p values 0.001 and 0.023 respectively) (Figure 3A and B and Supplementary Table 2). At high dose (4000IU), we found no significant effects of rs7041 (or other SNPs). Although higher doses may abrogate the genetic effect, better powered studies are needed to confirm this hypothesis.

**Figure 3:**
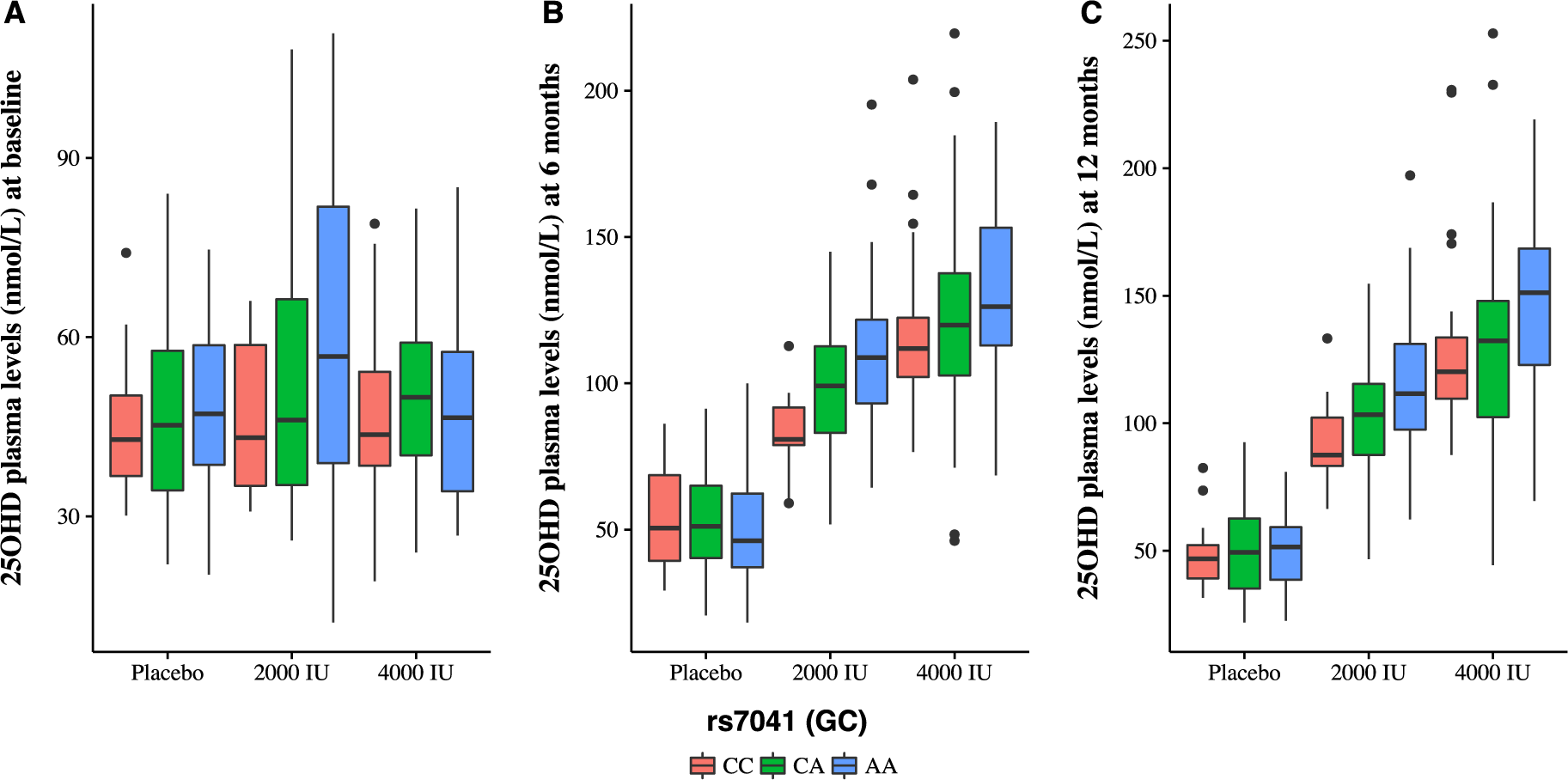
Genetic association analysis following vitamin D supplementation. Box-plots of rs7041 showing post-supplementation plasma levels of 25(OH)D by allocated treatment. Associations with rs7041 are statistically significant at both 6 (p=0.001, panel B) and 12 months (p=0.02, panel C) using permutation to derive empirical p-values after adjusting for baseline vitamin D levels and other relevant variables (see Methods and Supplementary Table 2). x-axis: genotypes, red (left) = CC, green (middle) = CA and blue (right) = AA. y-axis: 25(OH)D circulating levels (nmol/L) at baseline (A), 6 months (B) and 12 months (C).

We next considered whether there was evidence that genetic determinants of gene expression were modulated by supplementation with vitamin D by adopting a genetical genomics approach. Genetic variation is known to be an important determinant of individual gene expression and to be highly context-specific (13). We hypothesised that genetic variation may be an important contributor to individual gene expression differences in response to vitamin D. Although our study did not detect differences in gene expression after supplementation, we carried out an expression quantitative trait analysis as pre-specified given that individual level effects on gene expression dependent on genotype may still occur. Following sample processing and quality control, we analysed genotyping data on 497,136 variants for 288 individuals. We tested for evidence of association with gene expression using an additive linear model for 14,972 probes, including the top principal components as covariates after maximising for cis-eQTLs in each group (see Methods). We defined expression associated SNPs (eSNPs) as *cis*-eSNPs (those located within 1Mb of the gene expression probe) or *trans*-eSNPs (located >1Mb of the gene expression probe). At baseline, we found 31,568 cis-eQTLs and 34,254 at 12-months (unique SNP-probe pairs, 2000 IU and 4000 IU groups jointly to maximise sample size, FDR <5%) (Supplementary Table 3). There was a significant and positive overlap with comparable previously published data for whole blood eQTL (14) indicating high reproducibility despite large differences in sample size (1.75 fold-change overlap, q-value <0.001). To investigate response eQTLs present in samples from vitamin D supplemented individuals, we performed an analysis that took into account both effect size (gene expression differences) and statistical significance by obtaining eQTLs from the fold changes between treated (joint 2000 IU and 4000 IU 12 months' supplementation) and their baseline values after correcting for principal components as outlined above. We found no significant associations involving response eQTLs (FDR <10%, Supplementary Table 3, Supplementary Figure 8).

### Changes in plasma cytokine concentrations following vitamin D supplementation

In addition, we assessed whether supplementation with vitamin D had any significant effect on plasma levels of cytokines (Table 1). Consistent with lack of effect on gene expression results in the present and other previous studies (15), we did not identify any significant effects of vitamin D supplementation on plasma levels on cytokines. Likewise, we found no significant correlations between plasma levels of 25(OH)D and plasma levels of IFN-γ, IL-10, IL-8, IL-6 or TNF-α at 12 months (Supplementary Figures 9 and 10). Multivariate regression models testing the effect of supplementation after 12 months for either dose of vitamin D on changes in plasma cytokine levels (IFN-γ, IL-10, IL-8, IL-6 or TNF-α) after accounting for known confounders and baseline values did not show significant changes either (Figure 4 and Supplementary Table 4).

**Figure 4:**
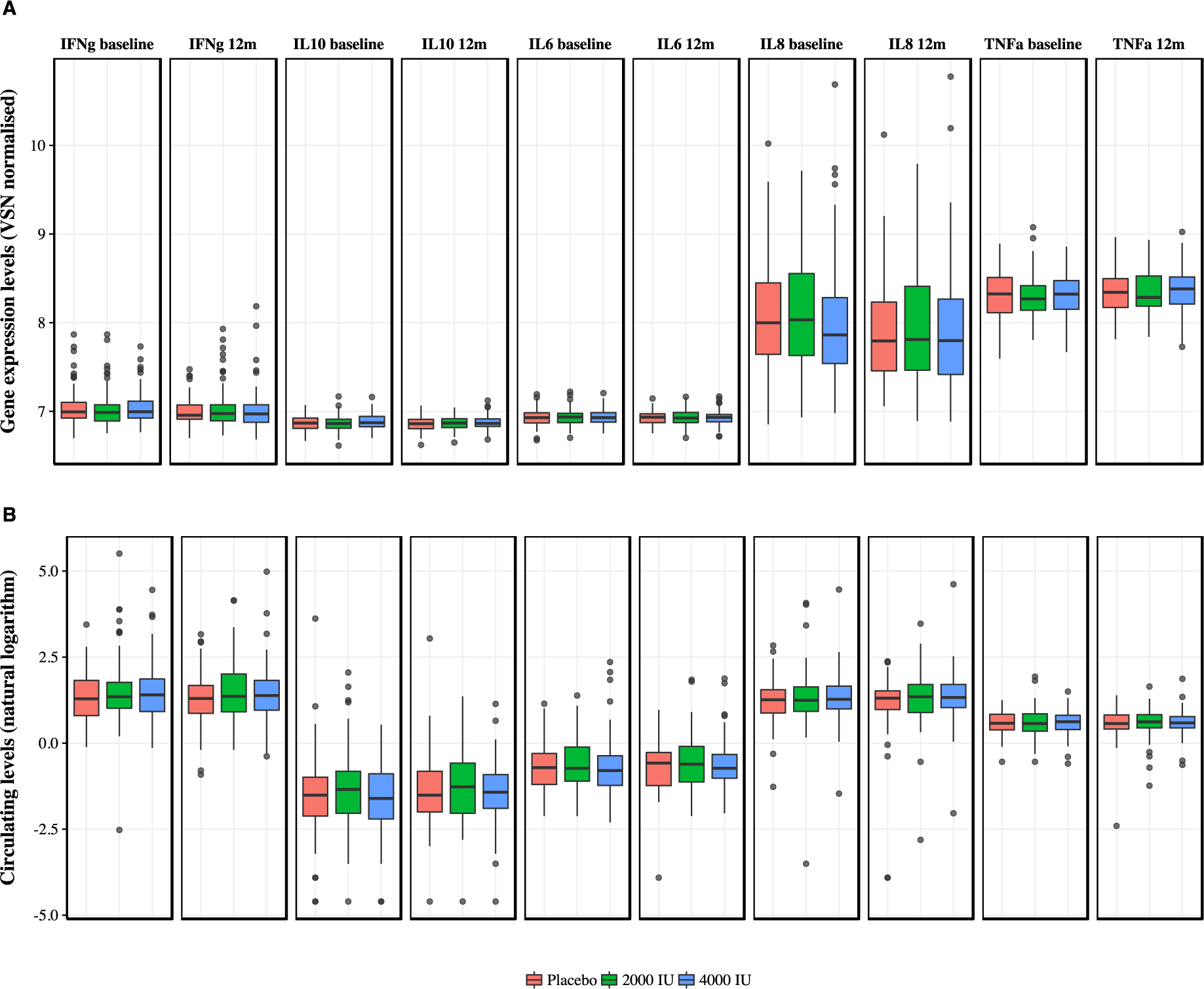
Boxplots of plasma cytokine levels and their corresponding transcripts before and after 12 months of vitamin D supplementation by allocated treatment. Gene expression (RNA) and circulating cytokine (protein) levels did not show changes after supplementation when accounting for baseline levels, known confounders and placebo (see Methods). VSN normalised mRNA levels (top row) and log-transformed protein levels (bottom row) of cytokines in whole-blood at baseline and 12 months (y-axis) for each trial arm (x-axis: red (left) = placebo, green (middle) = 2000 IU, blue (right) = 4000 IU).

## Discussion

Overall, the present study demonstrated that allocation to high-dose oral vitamin D_3_ in 305 older people had no significant effect on gene expression or plasma levels of cytokines when measured after 12 months despite achieving significantly higher plasma 25(OH)D levels (8). To our knowledge, this is the first randomised, placebo-controlled trial that assessed molecular changes and genetic effects following vitamin D supplementation.

Previous studies lacking appropriate randomization or use of placebo controls may have been confounded by time. Both seasonal and age related effects on gene expression have been reported previously (16, 17). Careful design and analysis are required; data analysis protocols can be pre-specified with blinding maintained. We demonstrated no significant effect of supplementation with either 2000 IU or 4000 IU of vitamin D on genome-wide gene expression or on plasma levels of IFN-γ, IL-10, IL-8, IL-6 and TNF-α after one year of supplementation. Although preliminary, the results of the present study suggest that plasma levels of vitamin D after supplementation can be modified by genetic variation.

Our study has several limitations. We did not collect samples within the first few hours or days following intervention and cannot exclude early transcriptomic changes. Indeed, other studies have observed changes in chromatin accessibility at candidate regions following vitamin D supplementation in peripheral blood mononuclear cells (18). Similarly, other tissues and specific cell types may show differences not observable in whole blood samples. We cannot address whether inflammatory processes deplete plasma levels of vitamin D, but our results do not support the hypothesis that long-term supplementation modulates plasma levels of cytokines. The study population were Caucasian, community-dwelling individuals. Younger individuals, those with particular diseases or different ethnicities may respond differently. Although we did not find differences for those with baseline vitamin D deficiency, our study was not designed to target this group.

This study demonstrated that genome-wide genetic variation, transcriptome and genetics of drug induced gene expression profiling can be integrated with biochemical and physiological measurements in the context of a randomised trial and may add important molecular and mechanistic insights. Interdisciplinary and collaborative efforts can guide the design and conduct of trials which require careful sample, data collection and analysis.

Further studies are needed to replicate the null associations and to assess why there were no long-term detectable differences after vitamin D treatment on gene expression. Physiological mechanisms that regulate the metabolism of vitamin D may achieve a steady state. Higher plasma levels of 25(OH)D may allow resources to be mobilised when needed. Upregulation of catabolic enzymes, unobserved confounding, and tissue specificity, amongst others, may account for the null findings. The present study highlights the difficulty in translating results from model organisms, in-vitro and ex-vivo studies and argues for further integration of molecular and clinical studies from in vivo observations.

## Materials and Methods

Details of the design, baseline characteristics and data analysis plan and results of the BEST-D trial have been previously reported (8, 9). Briefly, the primary objectives of BEST-D were to compare the effects on plasma levels of 25(OH)D and to determine the proportion of participants with plasma 25(OH)D levels >90 nmol/L after one year of supplementation with 4000 IU or 2000 IU of vitamin D_3_ versus placebo. Eligible participants were ≥65 years of age, living in the community and ambulatory. Participants were randomised in a ratio of 1:1:1 to each group using a minimisation algorithm balanced for age group (65–69, 70–74, ≥75 years), gender, body mass index (BMI), smoking history, ethnicity and history of fracture. Of 1122 individuals who were invited to participate, 313 (33%) agreed to a receive a visit from a study nurse for randomisation and 305 (32%) were successfully randomised between 24 September 2012 and 14 March 2013. All data and results were handled according to the trial and institutional guidelines in secure servers within the University of Oxford. All comparisons were conducted by intention-to-treat analyses using a pre-specified plan for the molecular data analysis (Supplementary Information).

### Procedures

Briefly, a research nurse visited participants at their homes to obtain medical history, samples and measurements. PAXgene Blood RNA tube (Qiagen) were used to ensure RNA stability without needing immediate processing. Biological samples were transported at 2 −4°C and then stored at −80°C. RNA and DNA samples were processed at the end of the study as detailed below. Plasma 25(OH)D levels were measured using an Access 2 immunoassay analyser (Beckman Coulter Ltd., High Wycombe, England) complying with the quality assurance DEQAS scheme. Further details have been described previously (8). Plasma cytokines were measured using a MesoScale Discovery multi-spot assay system. The V-Plex pro-inflammatory panel 1 kit was used, with detection antibodies for interferon-gamma (IFN-γ), interleukin 4 (IL-4), interleukin 6 (IL-6), interleukin 8 (IL-8), interleukin 10 (IL-10) and tumour necrosis factor alpha (TNF-α). In brief, assays were sandwich electrochemiluminescence (ECL) immunoassays. Between-run precision over the experiment for IFN-γ was 11.39% at 91.78 pg/mL and 10.73% at 10.73 pg/mL; for IL-4 12.99% at 11.35 pg/mL and 14.95% at 2.53 pg/mL; for IL-6 11.15% at 49.20 pg/mL and 13.02% at 8.85 pg/mL; for IL-8 9.38% at 32.67 pg/mL and 12.76% at 6.75 pg/mL; for IL-10 13.60% at 22.01 pg/mL and 13.82% at 4.49 pg/mL; and for TNF-α 12.29% at 11.15 pg/mL and 14.36% at 3.19 pg/mL.

#### DNA extraction and genotyping

Genomic DNA was extracted from the buffy coat layer using DNeasy Blood and Tissue Kit (Qiagen), and quantified by NanoDrop (Thermo Fisher Scientific; Waltham, MA) and Agilent 2100 Bioanalyzer (Agilent Technologies). 299 samples (of 305 possible) were available for DNA isolation and were processed over a single batch. Genotyping was performed using the Illumina Infinium HumanOmniExpress-24v1-0 (Illumina) beadchips following the Infinium HTS protocol (Illumina) at the Oxford Genomics Centre (WTCHG). Sample concentration was measured using PicoGreen (Thermo Fisher Scientific) and normalised. In total, 716,503 single nucleotide polymorphisms (SNPs) were genotyped. The genotype call rate cut-off was <98%. The overall call rate was 99.75% with one sample removed at this stage (genotype call rate 96.8%).

#### RNA extraction, cDNA conversion and microarray measurements

Total RNA was isolated from whole blood samples using PAXgene Blood RNA Kit (Qiagen) with recovery of RNA populations of <200 nucleotides and globin messenger RNA clearance using the GLOBINclear Kit (Ambion). Quantification of RNA and quality measures were assessed using Agilent 2100 Bioanalyzer (Agilent Technologies) and Nanodrop (Thermo Fisher Scientific). Complementary DNA synthesis, labelling and microarray hybridisation were performed at the Oxford Genomics Centre using Illumina Human-HT-12v4 expression BeadChip (Illumina). In total, 574 samples were available for further processing after microarray measurements (of 610 possible from the 305 participants who completed the study). Thirty samples were not available at end of study, four samples failed RNA quality metrics before array hybridisation and two during array processing. Three additional samples were identified as of low quality due to low amounts of complementary RNA upon further inspection. Arrays were run in batches of 96 randomised samples.

### Statistical analysis

#### A priori power for detecting gene expression differences

Statistical power calculations for gene expression analysis for *in vivo* studies are not well documented. We estimated statistical power using available data from vitamin D treated human cell line experiments (7, 19). We estimated that vitamin D_3_ supplementation may alter gene expression by 1.5 to 3-fold differences. Sample size was prioritised for the main trial outcomes (circulating levels of 25(OH)D). With a fixed sample size of 100 per group for the BEST-D trial, taking a two-sided alpha of 0.05 t-test, beta of 10%, standard deviation of 0.7, and equal size group we estimated we would be able to detect effect size differences of 1.25-fold change. Calculations were performed using the base package in R and pwr (v1.1-3).

#### Genome-wide SNP processing and statistical analysis

Quality control was carried out using standard approaches (20, 21) and included assessment of gender miss-identification; subject relatedness, duplication and divergent ancestry; individuals with elevated missing data rates or outlying heterozygosity rate; identification of markers (SNPs) with excessive missing data rates; identification of differing genotype call rates between groups; SNP quality (filtering of monomorphic SNPs, SNPs with missing values or nonsense values; low call rate; violation of Hardy-Weinberg equilibrium; duplication; and minimum allele frequency). We removed low quality markers followed by individuals. We excluded non-autosomal variants. We used the following criteria for filtering: call rates >98%; minor allele frequency (MAF) >10%; Hardy-Weinberg equilibrium threshold of 1×10^−6^. This resulted in 19 SNPs with a significantly different (p-value <0.01) missing data rate between cases and controls (treated vs placebo) being excluded; 4,893 variants due to missing genotype data; 20 variants due to Hardy-Weinberg equilibrium and 193,723 with MAF <10%. In total, 497,136 variants passed QC filters. Of 299 genotyped individuals, eleven were excluded after QC: one individual was excluded due to low genotype call rate (<98%); two due to gender misidentification; two due to relatedness (identity-by-descent value >0.1875); three due to ancestry other than Caucasian, and three due to a high genotype failure rate (≥0.03) and/or a heterozygosity rate ±3 SD from the mean (Supplementary Figure 1).

Linear regression association tests were conducted using frequentist methods with PLINK version 1.90 (22). We corrected for baseline vitamin D circulating levels, vitamin D intake (assessed at baseline), season (based on date of trial recruitment), gender, age, baseline BMI, medical history (incident fracture, incident respiratory infection, diabetes, heart disease, chronic obstructive pulmonary disease, asthma) and current smoking status. To explore genetic determinants of vitamin D levels we only considered SNPs previously identified by GWAS (10) and which were included in the genotyping array (rs12794714, rs2282679, rs7041 and rs7944926). We utilised an adaptive Monte Carlo permutation as implemented by PLINK to derive empirically determined significance values.

#### Quality control, normalisation of microarray data and differential gene expression analysis

Quality assessment of gene expression data included visual analysis of un-normalised data; analysis of built-in control probes; sample outlier detection and estimation of the proportion of probes expressed across samples (23) (Supplementary Figure 2). Outlier detection was carried out using ArrayQualityMetrics v3.24.0 (24). We removed samples that failed three criteria based on the package's internal scores of individual array quality, homogeneity between arrays and between array comparisons. We found that 11 samples were classed as outliers by all three methods. Outlier thresholds (sum of the distances to all other arrays, Kolmogorov-Smirnov statistic Ka and Hoeffding's statistic Da) were calculated by the package based on the array signal intensity values.

We excluded probes not expressed in at least three arrays with detection p-values <0.05. Pre-processing and probe filtering included background correction as described in (25) using built-in negative controls and VSN normalisation. A second procedure based on quantile normalisation (limma's neqc function) was used to test the main results of the differential expression analysis. We used the package illuminaHumanv4.db v1.26.0 as well as the manifest file for Illumina Human-HT-12v4 to annotate gene expression probes. We excluded probes from further analysis if probe sequences mapped to more than one genomic location; annealed at regions with SNPs present or mapped to non-autosomal locations. When mapping cytokines to their corresponding mRNA transcripts we found that only *IL-10* (ILMN_1674167) did not overlap known SNPs. Our final results filtered probes overlapping SNPs (Supplementary Table 1), but exclusion of these did not materially alter results. For comparison purposes, we present all cytokine transcripts (Figure 4). Principal components analysis was performed with R's prcomp function with scaling and centring.

We performed differential expression comparisons using limma v3.24.15 with linear models fit with empirical Bayes analysis (26). The primary comparison was a difference in difference estimator. We tested for linear or quadratic effects of vitamin D on expression and on the absolute change in expression. We also performed a linear mixed model analysis with the R package lmerTest (27), using person-specific random effects to account for between-person expression heterogeneity and fixed effects for time and time interacted with vitamin D levels.

#### Expression quantitative trait loci (eQTL) analysis

We used an additive linear model as implemented in the R package MatrixEQTL v2.1.1 (28) with inclusion of principal components (PC) from gene expression samples as covariates. We determined the number of PCs to correct for by running eQTL analyses with increasing numbers of PCs until the number of eQTL associations was maximised (29). Statistics and plots were carried out at the probe level. We used dbSNP human build 146, probe genomic locations as provided by Illumina, and p-value thresholds at <1e-8 for trans and <1e-5 for cis. Vitamin D response eQTLs were defined using fold change values without a cut-off threshold as input (instead of gene expression values) for association with SNPs (FDR <5%) after correcting for PCs per group. The maximum number of PCs to correct for was based on independent eQTL analyses of baseline and 12-months samples from gene expression values. We used R core packages, biglm (v. 0.9-1) and gvlma (v. 1.0.0.2) to regress PCs from gene expression values. Similar to the gene expression analysis, we tested for errors in the eQTL pipeline. Here we tested our results directly as comparable experiments have been done previously in whole blood samples (14). Despite differences in sample size, genotyping platforms and number of SNPs tested we chose a conservative genomic interval overlap test. We used the tool GAT (30) with the mappable genome as background and genomic intervals defined as plus and minus 1000 nucleotides for each SNP and ran the analysis of overlap between our results and (14) with 1000 permutations to obtain empirical p-values.

#### Statistical analysis of circulating cytokines

We imputed missing data using multiple imputation methods with 50 datasets, a maximum iteration of 50 and predictive mean matching (R packages mice v2.30 and miceadds v2.4-12) (31). No variable had more than 5% missing values. We performed analysis of covariance on each of the log natural transformed values of plasma levels of IFN-γ, IL-10, IL-8, IL-6 and TNF-α, accounting for the same confounders as in the genotype-25(OH)D analysis and including baseline values for every case. Linear regression summary tables presented were processed with the R package stargazer v2.3.1.

#### General software and plotting

R packages were run with R 3.2.4 (32). Custom scripts, sqlite3 (v. 3.13.0) and data.table (v. 1.9.6) were used for data processing. Figures were generated using package specific functions (limma and MatrixEQTL) or with ggplot2 (v. 2.1.0) and R's base plotting. Supplementary Figure 1 was plotted using code from (20).

## Main figures and tables

**Figure 1: CONSORT diagram for the functional genomics analysis of participants in the BEST-D study.** eQTL: expression quantitative trait loci.

**Figure 2: Global gene expression by allocated treatment with either vitamin D or placebo supplementation was assessed using principal components analysis of variance.** Principal components (PC) were calculated based on post-QC and VSN normalised gene expression signals. PC1 and PC2 (representing 15% and 8.5% of variance, respectively) are shown and coloured according to time-point (final visit at 12 months or at randomisation [baseline], panel A) and separately by treatment group (placebo, 2000 IU or 4000 IU, panel B).

**Figure 3: Genetic association analysis following vitamin D supplementation.** Boxplots of rs7041 showing post-supplementation plasma levels of 25(OH)D by allocated treatment. Associations with rs7041 are statistically significant at both 6 (p=0.001, panel B) and 12 months (p=0.02, panel C) using permutation to derive empirical p-values after adjusting for baseline vitamin D levels and other relevant variables (see Methods and Supplementary Table 2). x-axis: genotypes, red (left) = CC, green (middle) = CA and blue (right) = AA. y-axis: 25(OH)D circulating levels (nmol/L) at baseline (A), 6 months (B) and 12 months (C).

**Figure 4: Boxplots of plasma cytokine levels and their corresponding transcripts before and after 12 months of vitamin D supplementation by allocated treatment.** Gene expression (RNA) and circulating cytokine (protein) levels did not show changes after supplementation when accounting for baseline levels, known confounders and placebo (see Methods). VSN normalised mRNA levels (top row) and log-transformed protein levels (bottom row) of cytokines in whole-blood at baseline and 12 months (y-axis) for each trial arm (x-axis: red (left) = placebo, green (middle) = 2000 IU, blue (right) = 4000 IU).

Table 1: Selected characteristics at baseline and at 12 months by allocated treatment with vitamin D or placebo.

## Supplementary Information

Appendix 1: Data analysis protocol

Appendix 2: Lay summary

Supplementary Figure 1: Genotype data QC.

Supplementary Figure 2: Gene expression QC.

Supplementary Figures 3-6: Principal component analyses based on gene expression values.

Supplementary Figure 7: Gene expression p-value distributions.

Supplementary Figure 8: Quantile-quantile plot of fold change eQTLs.

Supplementary Figure 9: Heatmap of circulating cytokines and vitamin D.

Supplementary Figure 10: Relationship between circulating cytokines and vitamin D supplementation.

Supplementary Table 1: Differential gene expression comparisons in vitamin D trial. (A) Legend and description of all comparisons. (B) Main comparisons. (C) All comparisons (including those considered main [ST1B])

Supplementary Table 2: Genetic association results of candidate markers

Supplementary Table 3: Expression quantitative trait locus analysis

Supplementary Table 4: Summary of analysis of covariance of circulating cytokines

Supplementary Table 5: Vitamin D and Mendelian Randomization literature review

Supplementary Table 6: Literature review of vitamin D clinical trials in the 12 months to June 12, 2017

## Acknowledgements

We would like to thank all the trial participants, BEST-D Team, Wolfson Laboratory at CTSU and members of the Knight Lab and CGAT (<u>www.cgat.org</u>) for their support. AJBT was supported by the Medical Research Council (CGAT Fellowship), UK MED-BIO Programme Fellowship (MR/L01632X/1), the Multiple Sclerosis Society UK (Grant 915/09) and the Council for Science and Technology (CONACyT, Mexico, Grant 211990). JK was supported by NIHR Oxford Biomedical Research Centre, the European Research Council under the European Union's Seventh Framework Programme (FP7/2007-2013) (ERC Grant agreement no. 281824) and Wellcome Trust Investigator Award (204969/Z/16/Z). The Clinical Trial Service Unit and Epidemiological Studies Unit (CTSU) at the University of Oxford received funding from the UK Medical Research Council, the British Heart Foundation and Cancer Research UK. We thank the High-Throughput Genomics Group at the Wellcome Trust Centre for Human Genetics (Funded by Wellcome Trust grant reference is 090532/Z/09/Z and MRC Hub grant G0900747 91070) for the generation of data. **Role of the funding source:** The study received funding from the British Heart Foundation, UK Medical Research Council and CTSU, University of Oxford. The British Heart Foundation (PG/12/32/29544) and British Heart Foundation Centre for Research Excellence provided partial funding for the study. Active and placebo vitamin D capsules were kindly donated by Tischcon Corporation (Westbury, New York, USA). The funders had no role in data collection, analysis, interpretation or writing of the report. All authors had access to all the data in the study. Trial registration: SRCTN Number 07034656; EudraCT Number 2011-005763-24.

## Author contributions

AJBT and JCK conceived the study. AJBT, JA, RC and JCK designed the study. JA and RC are the principal investigators of the BEST-D trial. AJBT performed the analysis with input from AD, DS, AH and JE. KP, EL and MH performed experiments. JA, RC and JCK provided senior supervision. AJBT and JCK wrote the manuscript with contributions from all authors. **Competing interests**: None. **Data and materials availability**: Gene expression data are available through ArrayExpress (E-MTAB-6246). Cytokine, phenotype and genotyping data are available from CTSU, University of Oxford through a material transfer agreement prior consent. All computational code used for processing and analysis is available at <u>https://github.com/AntonioJBT/BEST_D</u>.

